# Detecting adaptive changes in gene copy number distribution accompanying the human out-of-Africa expansion

**DOI:** 10.1101/2023.08.14.553171

**Authors:** Moritz Otto, Yichen Zheng, Paul Grablowitz, Thomas Wiehe

## Abstract

Genes with multiple copies are likely to be maintained by stabilizing selection that puts a bound to unlimited expansion of copy number. We designed a model where copy number variation is generated by unequal recombination, which fits well to a number of genes, surveyed in three human populations. Based on this theoretical model and on computer simulations, we were interested in answering the question whether gene copy number distribution in the derived European and Asian populations can be explained by a purely demographic scenario or whether shifts in the distribution are signatures of adaptation. Although copy number distribution in most of the analyzed gene clusters can be explained by a bottleneck as in the out of Africa expansion of homo sapiens 60-10kyrs ago, we identified several candidate genes, for instance `AMY1A` and `PGA3`, whose copy numbers are likely to be selected differently among African, Asian and European populations.

## Introduction

Gene copy number variation (CNV) refers to the presence of multiple copies of a gene family within a genome, resulting from duplications, deletions, or rearrangements. Combined with their high mutation rate CNVs constitute a significant driver of genomic variability that allows for rapid adaptive evolution in response to environmental changes (Sudmant et al. 2015; Brahmachary *et al*. 2014; Carvalho and Lupski 2016; Iskow et al. 2012; Sebat et al. 2004).

A well studied example of CNV within human population is provided by the salivary amylase gene, whose variations in the number of copies are hypothesized to correlate with the extent of dietary starch consumption not only in human but also in other species (Pajic et al. 2019; Atkinson et al. 2018; Carpenter et al. 2015; Usher et al. 2015; Falchi et al. 2014; Perry et al. 2007).

In general, copy number variation may result from different evolutionary forces acting upon them. Demographic events, such as population migrations and expansions, can lead to changes in gene frequencies and distributions over time. Simultaneously, natural selection acts on genetic variations, favoring advantageous alleles and promoting their proliferation within populations.

It is known that both demographic effects and selection may produce similar patterns in single nucleotide as well as in structural variants, making it difficult to disentangle these forces (Lohmueller 2014; Stajich 2004). For SNP or allele frequency data, there have been well-developed statistics (e.g., (Tajima 1989; Fu 1997)) that are “standardized” so that a genomic baseline can be established, from which loci under selection may be detected. However, such a genomic baseline is not available for gene copy number variation data. Therefore, we resorted to a more basic approach involving modelling and computer simulations.

We have recently examined the evolutionary dynamics of multi-copy gene families with respect to selective pressure and unequal recombination (Otto *et al*. 2022). This study focused on analyzing the impact of stabilizing selection on gene copy numbers, while considering the role of recombination as a randomizing mechanism that introduces variability within the population.

By expanding this model, we aimed to assess whether gene copy number alterations observed within human populations could be solely attributed to demographic events or whether selective pressures have played a role in shaping these variations.

In this study, we conducted extensive simulations under various scenarios of human demography and selective changes. By disentangling the effects of these two forces, we sought to gain a deeper understanding of the evolutionary processes driving gene copy number variation in human populations. Based on empirical data of human gene copy numbers we identified several candidate genes, whose copy numbers are likely to be selected differently among African, Asian and European populations.

## Materials and Methods

### Gene copy number variation in human

We started with the dataset provided by Brahmachary *et al*. (2014). Using Nanostring technology they estimated gene copy numbers of 180 gene-families in 165 individuals of three populations (60 African Yuroba - YRI, 60 Central Europe - CEU and 45 Asia - CHB) based on data collected in the framework of the 1,000 Genomes Project (Sudmant *et al*. 2010).

While some of these loci showed copy numbers of *>* 100 copies (DUX4 even up to 600), we focused on intermediate copy numbers and removed all satellite loci, genes on sex chromosomes, genes with minimum copy number below 2, and genes with mean copy number (in YRI) below 5 or above 60. For genes that have two primer sets, only one is taken. We used *t*-test and *F*-test statistics to select gene families with significant differences in mean or standard deviation between YRI-CHB or YRI-CEU comparisons and removed those that showed no statistical evidence in any of these. This resulted in 42 gene families, see Table 1 and the copy number distributions of four of them are shown in Fig 1.

**Table 1.**
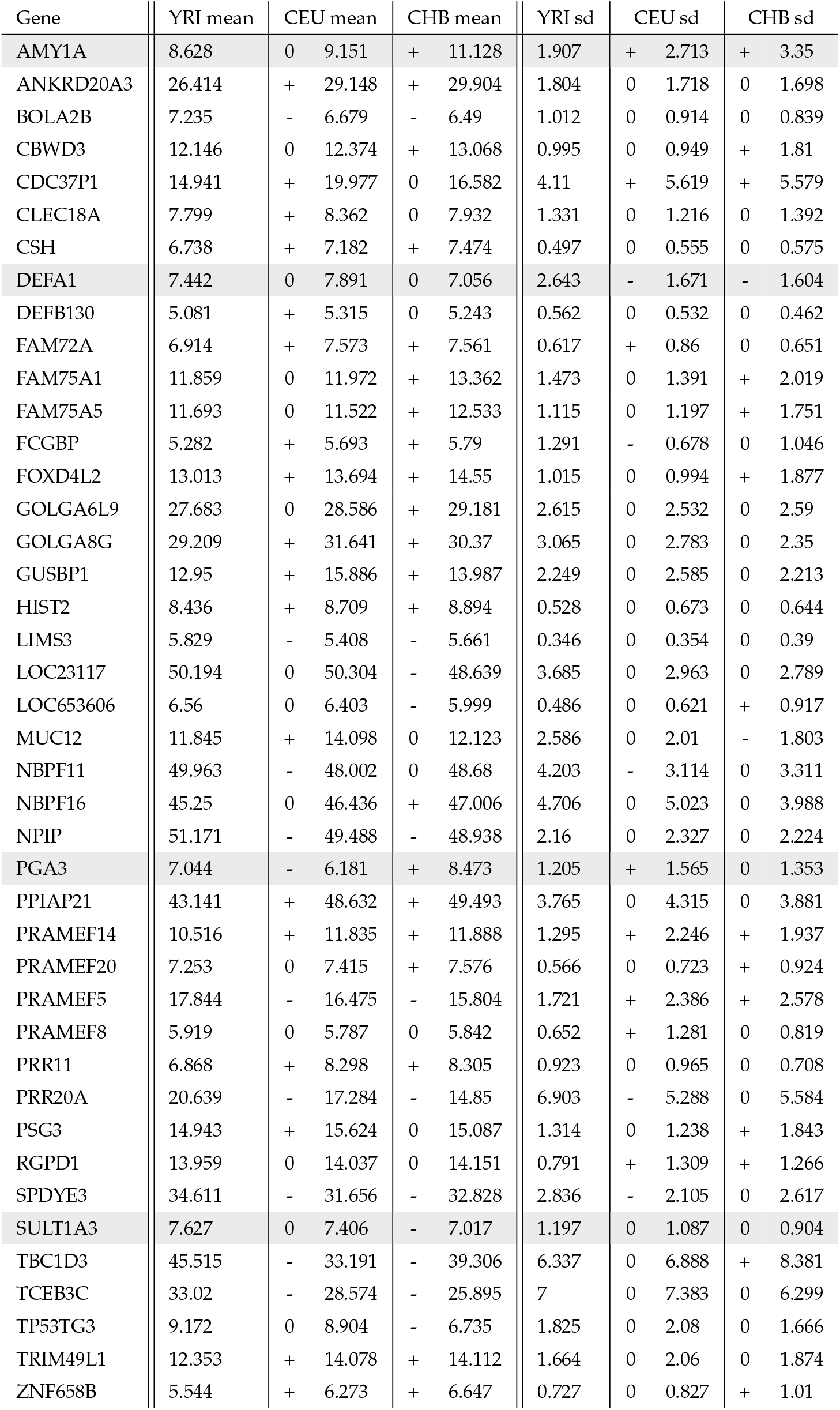
Differences in mean and variance between populations. 0 indicates no significant change, ‘+’ a significant increase and ‘–’ a significant decrease (*t*-test for the mean and *F*-test for the standard deviation; *α* = 0.05). The four candidate genes shown in Fig 1 are highlighted with lightgray background.

**Figure 1.**
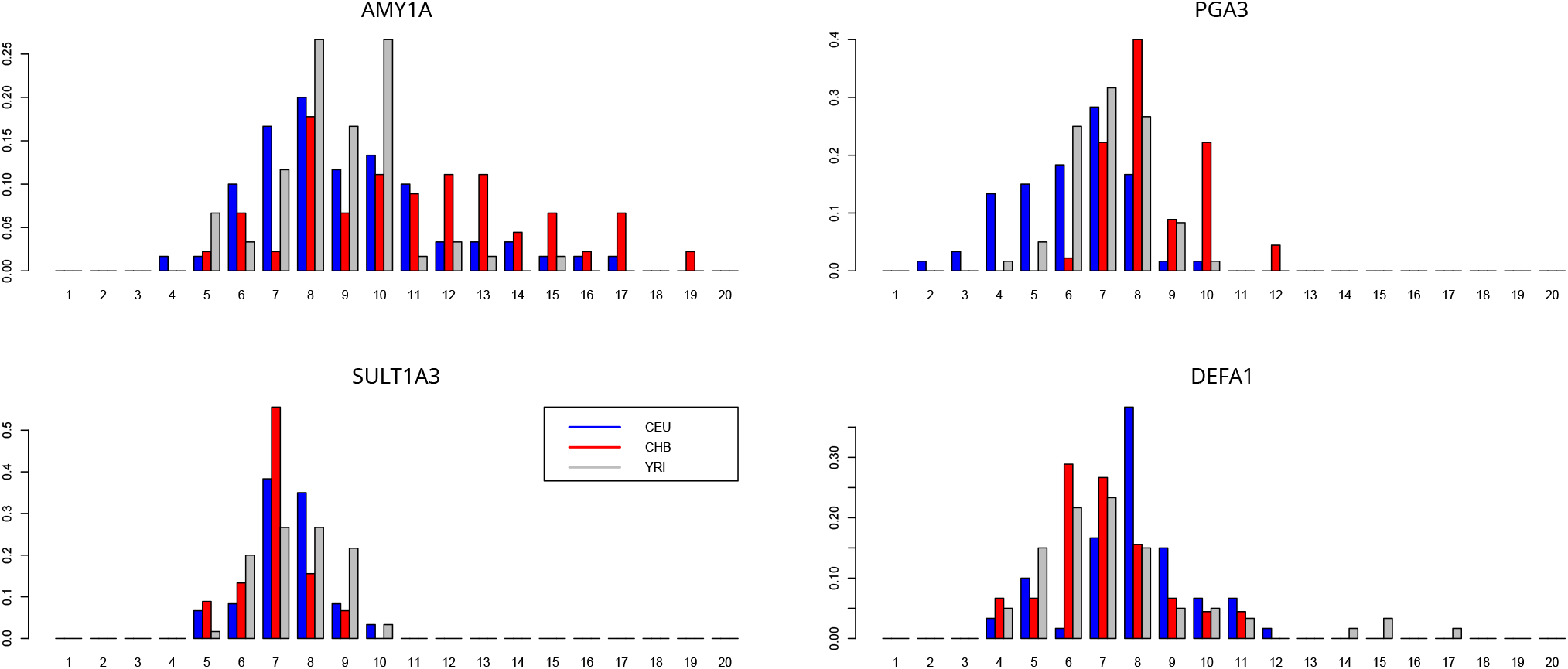
Gene copy number distribution in four exemplary gene famiies in three human populations, CEU, CHB, YRI. Data adapted from (Brahmachary *et al*. 2014).

### Unequal recombination model

In a recently developed model we considered unequal recombination and selection to describe the evolution of tandem gene arrays (Otto et al. 2022). We shortly summarize the main findings. Consider two chromosomes with gene arrays of size *y*_1_ and *y*_2_. A recombination event happens at rate *r* and may produce a gamete of gene array size according to the trapezodial distribution, such that

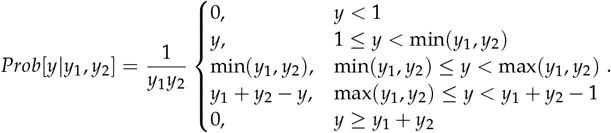

See Fig 2A for an illustration. We apply a fitness function, where each newly arising copy has a positive, yet decreasing benefit *s*_*x*_. This is motivated by assuming a beneficial effect, yet with diminishing returns, either of increased gene dosage or of increased allelic diversity within an individual (Otto et al. 2022). At the same time, we assume additional copies to be selected against with an increasing selective disadvantage *s*_*y*_. This is motivated by assuming an increasing cost of replication, of gene processing and of maintaining genome integrity. Both effects are cast in a double-epistatic fitness function with two selection coefficients (*s*_*x*_, *s*_*y*_), governed by a single epistasis parameter (*ε*). To avoid the trivial long-term evolution equilibrium of one copy, we assume *s*_*x*_ *> s*_*y*_.

**Figure 2.**
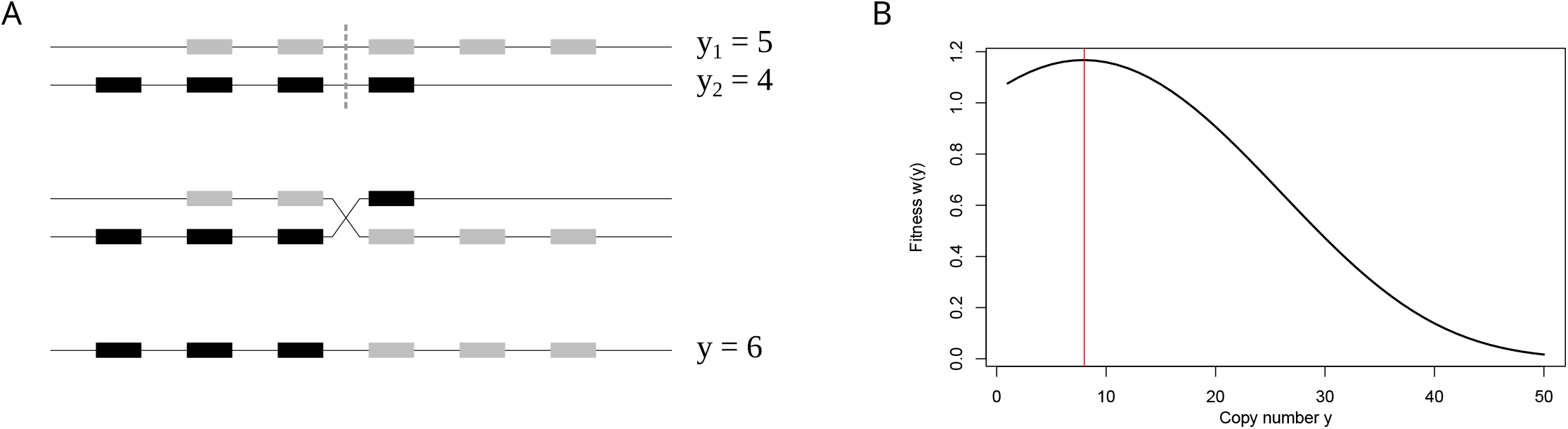
A. Sketch of the unequal recombination process. Starting with two chromosomes with *y*_1_ = 5 and *y*_2_ = 4 gene copies, two break points are chosen. One of the recombinants is then propagated. Its copy number (here *y* = 6) is Trapezoidal, as shown in (Otto *et al*. 2022). B. Example of the fitness function *ω*(*y*) (equation (1)) with *ε* = 0.05, *s*_*x*_ = 0.05, *s*_*y*_ = 0.0025, which leads to an optimal copy number *y*_opt_ *≈* 8 copies.

Furthermore, we assume *ε* = 0.05 to be constant in the following. Summarizing, fitness of a diploid individual with total gene copy number *y* is given by

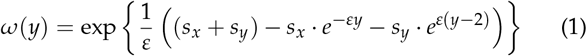

This leads to an optimal copy number *y*_opt_ of

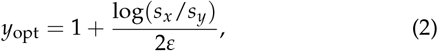

which is determined by the ratio *s*_*x*_ /*s*_*y*_ when *ε* is kept fixed. See Fig 2B for an example. The population is then simulated according to a Wright-Fisher model with non-overlapping generations and with selection and recombination described as above. It was shown, that in the deterministic model the equilibrium copy number distribution is centered around *y*_opt_ and is well approximated by a Gamma distribution (Otto *et al*. 2022). Furthermore, the co-efficient of variation 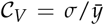 is correlated to the logarithm of the recombination - selection ratio log(*r*/*s*_*x*_). With strong selection and low recombination the distribution is tightly distributed around the optimal value, whereas higher *r* and lower *s*_*x*_ lead to a widespread distribution. For convenience, we introduce two new parameters:

- *q*_*S*_ = *s*_*x*_ /*s*_*y*_, the *’selection ratio’*, which determines the optimal copy number, such that for *ε* = 0.05 we find

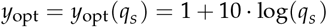
- *q*_*R*_ = *r*/*s*_*x*_, the *’recombination/selection ratio’*, which measures the impact of unequal recombination compared to the selective pressure of the fitness function and therefore determines the coefficient of variation 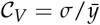 of the equilibrium distribution.

### Regression

We aim to quantify the effect of (*r, s*_*x*_, *s*_*y*_) on the resulting equilibrium copy number distribution and – vice versa – to estimate for given empirical data the underlying parameter triple. To analyze the equilibrium distribution of the unequal recombination process under drift, we simulated the population evolution under different parameter settings. The codes for all following simulations are available at https://github.com/y-zheng/Distinguishing-roles-adaptation-demography-gene-copy-number-changes-human-populations. Population size is kept at *N* = 5, 000 and assumed to be at an initial state of 5 copies on each chromosome. The different input parameters are given in Table 2.

**Table 2.**
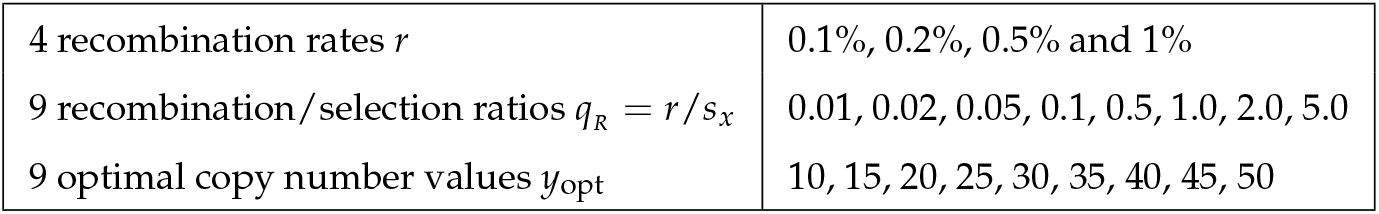
Parameters for regression simulations.

Together, they define 324 triples *r, s*_*x*_, *s*_*y*_. Additionally, we generated 160 random pairs such that *q*_*R*_ is between 0.01 and 5 and *y*_*opt*_ is between 4 and 60 and combined them with the four recombination rates, leading to a total parameter set of 964 combinations, where we disregarded those triples with selective strengths *s*_*x*_ *>* 0.1 to keep a realistic parameter range.

For each of this parameter combinations, we evolve the population under the given selection scheme for 5 million generations. The first 200,000 generations were discarded as burn-in and the population statistics (mean copy number 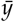 and standard deviation *σ*) are recorded every 20,000 generations.

In total, this results in *≈* 185, 000 data points, which we used to determine the relationship of input parameters (*r, s*_*x*_, *s*_*y*_) and output population statistics 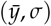.

As indicated in (Otto et al. 2022) we suggest a mean copy number 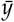 close to its optimal value *y*_opt_ and a correlation of *𝒞*_*V*_ to log(*q*_*R*_). Indeed, with *r*^2^-values of 0.9842 and 0.9088 we find

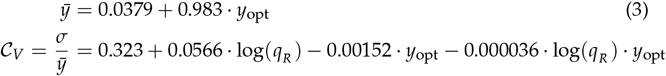

We calculated the *q*_*S*_ and *q*_*R*_ ratios based on 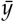 and 𝒞_*V*_ from gene copy numbers (see Table 1) using the regression formula (3) with four recombination rates *r* = 0.001, 0.002, 0.005 and 0.01. Results for the four candidate genes shown in Fig 1 are given in Table 3.

**Table 3.** Estimates of selection coefficients *s*_*x*_, *s*_*y*_ under four recombination rates *r* = 0.001, …, 0.01 based on regression equation (3). The displayed gene families are the ones of Fig 1 for all three populations. Values in parentheses are out of the range 0.001 *< s*_*x*_ *<*?0.1 in YRI and hence not used in simulations.

| Gene | Pop | Mean | SD | $y_{opt}$ | $r = 0.001$ | | $r = 0.002$ | | $r = 0.005$ | | $r = 0.01$ | |
| --- | --- | --- | --- | --- | --- | --- | --- | --- | --- | --- | --- | --- |
| | | | | | $s_x$ | $s_y$ | $s_x$ | $s_y$ | $s_x$ | $s_y$ | $s_x$ | $s_y$ |
| AMY1A | CEU | 9.1511 | 2.7133 | 9.2708 | 0.0012 | 0.0005 | 0.0025 | 0.0011 | 0.0062 | 0.0027 | 0.0125 | 0.0055 |
|  | CHB | 11.128 | 3.3503 | 11.282 | 0.0011 | 0.0004 | 0.0022 | 0.0008 | 0.0054 | 0.0019 | 0.0109 | 0.0039 |
|  | YRI | 8.6279 | 1.9074 | 8.7386 | 0.0048 | 0.0022 | 0.0097 | 0.0045 | 0.0242 | 0.0111 | 0.0483 | 0.0223 |
| PGA3 | CEU | 6.1808 | 1.5646 | 6.2491 | 0.0029 | 0.0017 | 0.0058 | 0.0035 | 0.0146 | 0.0086 | (0.0292) | (0.0173) |
|  | CHB | 8.4731 | 1.3526 | 8.5811 | 0.0144 | 0.0068 | 0.0289 | 0.0135 | 0.0722 | 0.0338 | (0.1445) | (0.0677) |
|  | YRI | 7.0444 | 1.2053 | 7.1277 | 0.0122 | 0.0066 | 0.0245 | 0.0133 | 0.0611 | 0.0331 | (0.1223) | (0.0663) |
| SULT1A3 | CEU | 7.4058 | 1.0872 | 7.4953 | 0.0186 | 0.0097 | 0.0373 | 0.0195 | 0.0932 | 0.0487 | (0.1865) | (0.0974) |
|  | CHB | 7.0165 | 0.9041 | 7.0993 | 0.0259 | 0.0141 | 0.0518 | 0.0282 | 0.1295 | 0.0704 | (0.2591) | (0.1408) |
|  | YRI | 7.6269 | 1.1971 | 7.7202 | 0.0155 | 0.0079 | 0.0311 | 0.0158 | 0.0774 | 0.0395 | (0.1548) | (0.0791) |
| DEFA1 | CEU | 7.8911 | 1.6708 | 7.9889 | (0.0058) | (0.0029) | (0.0116) | (0.0058) | 0.0291 | 0.0145 | 0.0581 | 0.0289 |
|  | CHB | 7.0561 | 1.6041 | 7.1396 | (0.0045) | (0.0024) | (0.0091) | (0.0049) | 0.0225 | 0.0122 | 0.0451 | 0.0244 |
|  | YRI | 7.4421 | 2.6428 | 7.5321 | (0.0005) | (0.0002) | (0.0009) | (0.0005) | 0.0023 | 0.0012 | 0.0046 | 0.0024 |

### Demography simulations

To determine whether significant changes of mean and variance of the copy number distribution (Table 1) can be explained by demographic history of human populations, we examined in total 6 different scenarios (enumerated as I - VI), see Fig 3.

**Figure 3.**
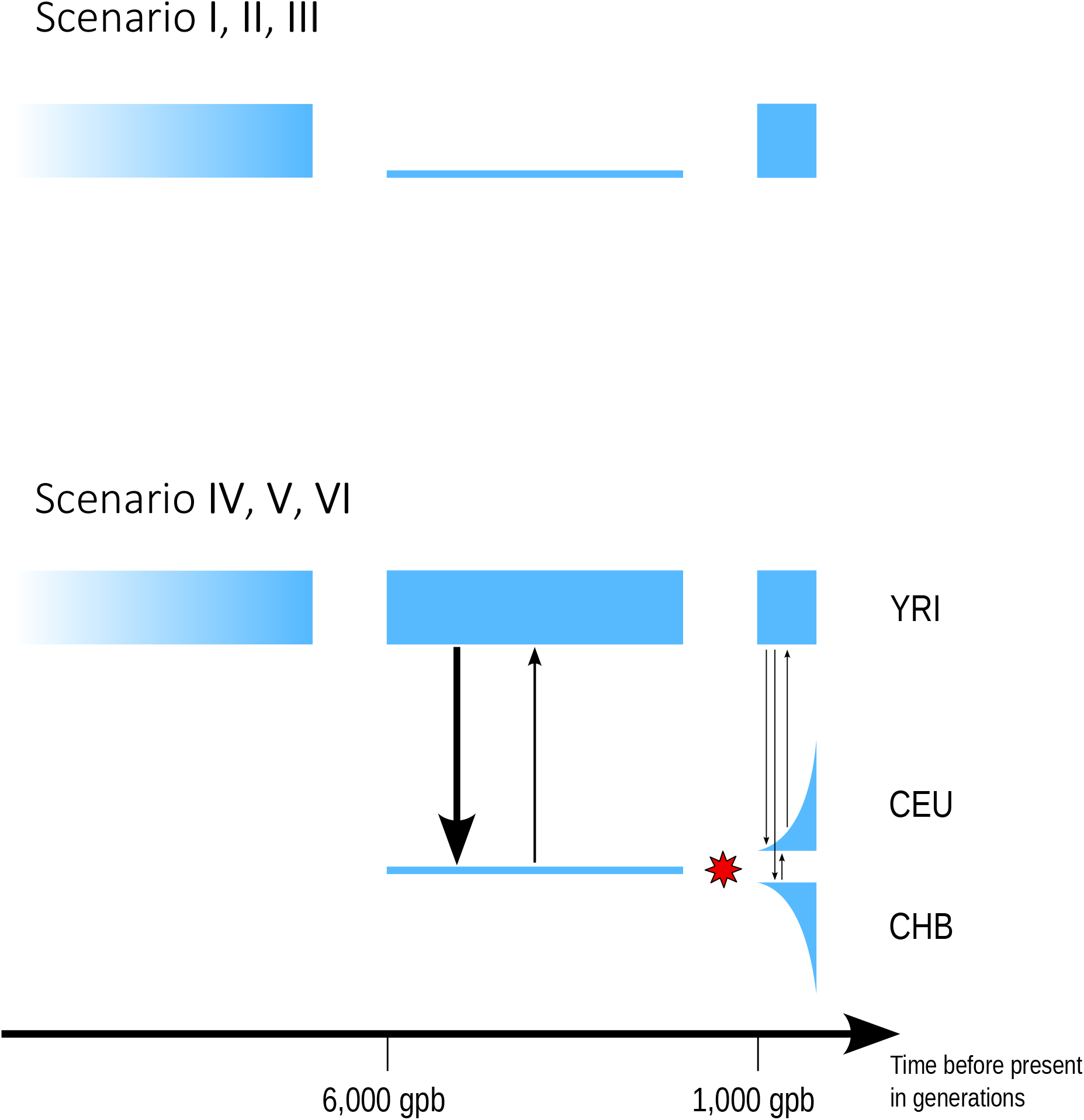
Illustration of six different scenarios investigated. Scenarios I-III: simple bottleneck lasting 5,000 generations. Reduction to 1% (scenario I), 5% (II) and 10% (III) of its original size (N=10,000). Scenarios IV-VI: GADMA model of human demographic history. Without change of selection intensity (scenario IV), with change of selection intensity (symbolized by the red star) 896 generations ago (scenario V) and 500 generations ago (VI). Black arrows indicate direction and frequency of migration between subpopulations.

#### Simulation of the bottleneck model

First, we ran a simple bottleneck model of three different population reductions. Each is divided into three phases: (1) Burn-in phase. For each gene we used the estimated (*r, s*_*x*_, *s*_*y*_)-triple based on the dataset from YRI. These parameters were chosen as input to produce an equilibrium population of *N* = 10, 000 by a burn-in process of 200,000 generations. ‘Independent’ equilibrium populations are produced by recording the population state every 20,000 generations. (2) Bottleneck. From equilibrium we reduced the population size to *N* = 100, 500 or 1, 000, denoted scenario I, II and III, and kept it such for 5,000 generations. (3) Recovery phase. At the end of the bottleneck, the population is reset to *N* = 10, 000 and the copy number distribution is recorded every 50 generations until generation 1000 after the bottleneck. We ran the bottleneck simulations I - III on all gene families given in Table 1, with recombination rates *r* = 0.001, 0.002, 0.005 and 0.01, and discarded parameter combinations with *s*_*x*_ outside the interval [0.001, 0.1] in YRI. This gives a final total of 42 gene families and 95 gene-*r* combinations. For each gene, recombination rate and bottleneck population size combinations, 10,000 replicates are produced (from 100 ‘independent’ starting equilibria).

We then traced mean and 𝒞_*V*_ along the recovery phase and compared these with the empirical data from CHB and CEU populations.

#### Simulation of the human population history

A more realistic population history of human is given by the *Genetic Algorithm for Demographic Model Analysis* (GADMA) (Noskova *et al*. 2020), which also includes migration between subpopulations. We ran simulations on four candidate genes (`AMY1A, PGA3, SULT1A3, DEFA1`) with the following modification of the GADMA-demography: As ancestral population (*N* = 9, 900 in GADMA), we used the equilibrium populations (*N* = 10, 000) from the previous section. Therefore, we started the simulation 5992 generations before present, roughly corresponding to the onset of the ‘out-of-Africa’ expansion, when the Eurasian population split from the ancestral population and experienced a sharp bottleneck. To reduce computation time, we did not simulate the continued evolution of the African (YRI) population, since we assumed it to be in equilibrium; for migration from YRI to Eurasian populations, we drew samples from the ancestral population. At 896 generations before present, CEU and CHB split from each other and started to evolve including reciprocal migration and exponentially increasing population size. In the following, we refer to this simulation as scenario IV. At ‘present’, copy number distributions (mean and variance) were recorded. For each gene and recombination rate combination, 10,000 replicates were produced.

We also ran the same population model with a change of the selection parameter either at 500 generations or 896 generations before present (the latter being the CEU/CHB split time). The new selection parameters (*s*_*x*_ and *s*_*y*_) are different for CEU and CHB populations, and are estimated from present CEU/CHB distributions (see Table 3). These simulations are hereafter called scenario V (selection change 896 generations before present) and VI (500 gpb).

### Rejecting a purely demographic model

Observing the copy number distribution for a gene family in the ancestral (YRI) population, we seek to answer the question whether the observed distribution in the derived population (CEU or CHB) can be explained by a purely demographic model (various bottlenecks, but keeping selection pressure constant as modelled in scenarios I to IV) or not (demography plus change in selection pressure as modelled in scenarios V or VI). To decide this we use the following strategy. For each scenario I-VI and each parameter triple estimated from the YRI population, 10,000 replicates were produced. From each resulting equilibrium distribution, we record the mean 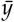 and standard deviation *σ*. This results in a parameter distribution for each scenario. If a chosen empirical data set has mean 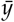 or standard deviation *σ* which are not in the 95% quantile of the 10,000 simulated values, we conclude that this particular scenario is to be rejected as a possible explanation of the data. In particular, we reject a purely demographic explanation, if scenarios I-IV are rejected.

## Results and Discussion

In this study, we conducted an analysis of multicopy gene family evolution using a model that incorporates unequal recombination and selection. Our investigation aimed to examine the copy number changes observed in subpopulations of Europe, Asia, and Africa and to determine whether these changes could be attributed to either constant selective pressure and demographic factors or an adaptive change of selection together with demography. Our findings reveal that the observed copy number variations in several genes cannot adequately be explained by demographic processes alone, suggesting a – possibly adaptive – change of selective pressure in the derived populations.

Based on the data of Brahmachary *et al*. (2014), we chose 42 gene families of intermediate copy number that show significant differences in their distribution among different populations (Table 1). Although the raw data rely on phase I of the 1000 Genomes project they proved to be most suitable for our analyses. More recent data, for instance from the human pangenome project (Liao *et al*. 2023), still lack sufficient coverage of the different subpopulations.

When comparing the copy number distributions of the 42 candidate genes in the Asian and European with the African population (assumed to be in equilibrium), we observe 61 significant changes in mean copy number and 29 significant changes in the variance (Table 1), of which only seven show a decrease of variance (one example is `DEFA1`). Within our model, assuming constant recombination rate among subpopulations, and no demographic changes, a decreased variance (or standard deviation) can only be achieved by an increase of positive selection (*s*_*x*_), since 𝒞_*V*_ is determined by *q*_*R*_ = *r*/*s*_*x*_, see equation (3). However, the most common case is that of a consistent significant shift in the mean in both derived populations without affecting the variance, i.e. either (+ + I 00) or (- - I 00), which occurs in 12 of the 42 analyzed genes. Only one gene (`PGA3`) showed opposite significant changes of the mean (increasing in Asia but decreasing in Europe).

To test whether a change of population size is sufficient to explain these significant differences in copy number statistics (shown in Table 1), we ran the simple bottleneck scenarios I-III with 95 parameter combinations (*r, s*_*x*_, *s*_*y*_) based on the ancestral copy number distribution of YRI and the regression equation (3). As an example, for PGA3 and *r* = 0.001 we ran simulations with selection coefficients *s*_*x*_ = 0.0122 and *s*_*y*_ = 0.0066, see Table 3. Fig 4 shows mean gene copy number of 10,000 simulated bottleneck populations over time for each recombination strength (*r* = 0.001, 0.002 and 0.005). Note that for this gene the value *r* = 0.01 was neglected, since the selection strength *s*_*x*_ would exceed the threshold of 0.1. Gray boxes indicate the centered 50% quantile, white the 95% and whiskers the 99% quantiles. With strong bottleneck (reduction to *N* = 100 for 5,000 generations) and under low recombination and, hence, weak selection (*r* = 0.001, and *q*_*R*_ = *r*/*s*_*x*_, *q*_*S*_ = *s*_*x*_ /*s*_*y*_ constant) we find the widest variation among the 10,000 replicates. Higher *r* and stronger selection result in a mean value close to the one of YRI population, i.e. the value we would expect from the initial parameter estimation.

**Figure 4.**
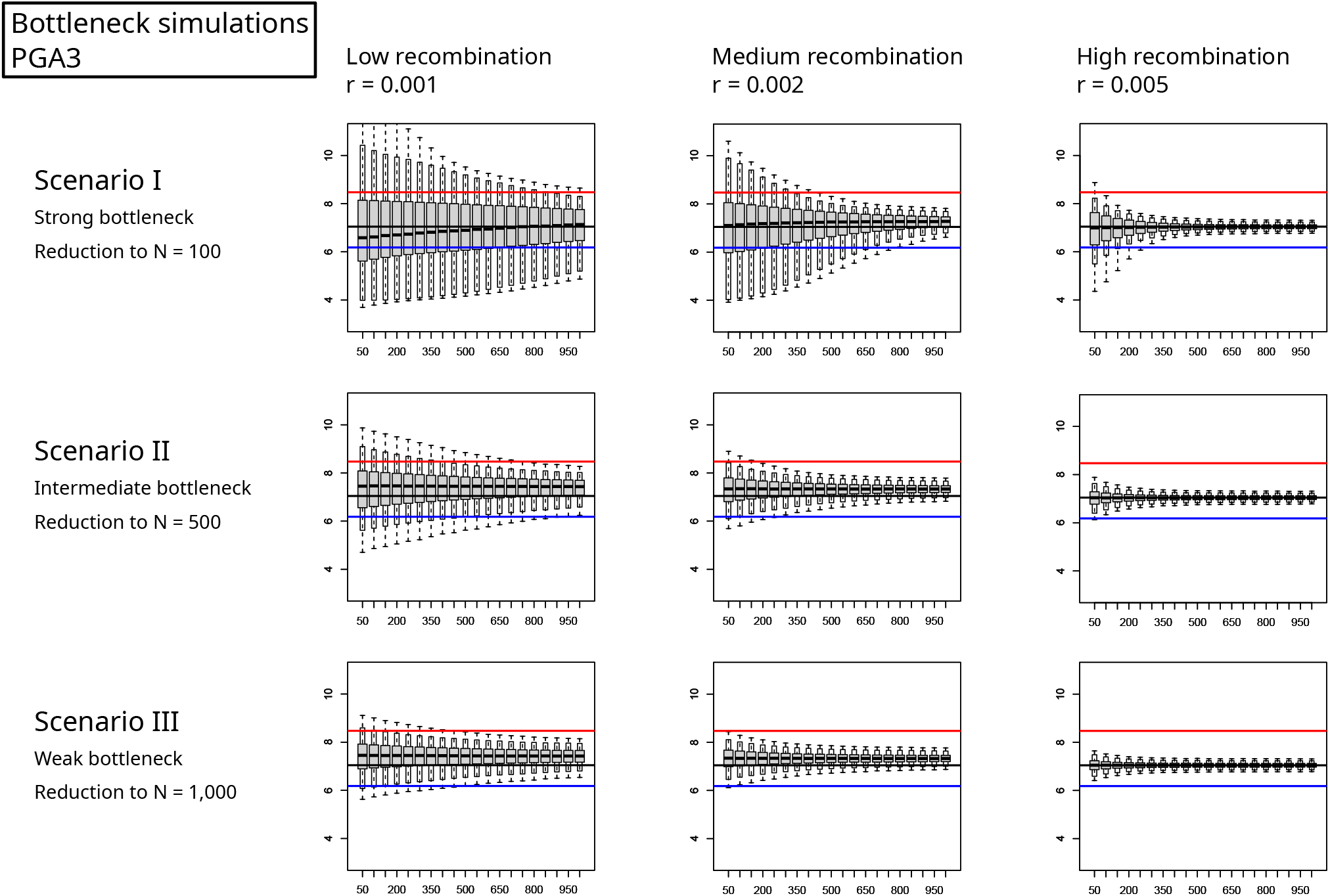
Mean copy number over time. After population reduction to *N* = 100, 500, 1000 (top to bottom) we traced the mean-value of 10,000 replicates over time (x-axis in generations). The input parameters *s*_*x*_, *s*_*y*_ were estimated for *r* = 0.001, 0.002, 0.005 (left to right) from the YRI-data set for the candidate gene `PGA3` (see Table 3) and kept constant over time, to see the effect of the bottleneck and recovery. Whiskers mark the 99% quantile, the white box the 95% quantile. Horizonal lines mark the values from the original data set of Brahmachary *et al*. (2014) (black: YRI, red: CHB, blue: CEU).

In this particular example, the empirical data show a signifcantly higher mean copy number of `PGA3` in CHB (red line) and a lower mean value in CEU (blue line) compared to YRI (black horizontal line). It is the only gene in our set, that shows significant shifts of mean copy number in opposite directions. Only under a strong bottleneck and with low recombination these changes could be explained without invoking a change of selection intensity.

The results for all 95 parameter combinations obtained for scenario I (strong bottleneck) are summarized in Table 4. To test whether observed means or standard deviations can be explained by scenario I, we considered the time point after 1,000 generations of recovery (first row of Fig 4, last boxplot in each panel). If the mean or resp. 𝒞_*V*_ lies within the 95% quantile, we indicate non-significant differences with a 0. Significant changes are marked with a single *\** (*α* = 5%) or double asterisk *\* \** (*α* = 1%). Taking again `PGA3` as example, we find a mean value which is significantly smaller in CEU than in YRI (marked with –). With *r* = 0.001, this might be explained by a bottleneck (denoted by 0), whereas for *r* = 0.002 and *r* = 0.005 we find a significant difference (*\*\**) and the bottleneck explanation to be highly unlikely. Higher recombination *r* = 0.01 led to *s*_*x*_ values greater than 0.1 in CHB and YRI (see Table 3) and hence was omitted.

**Table 4.**
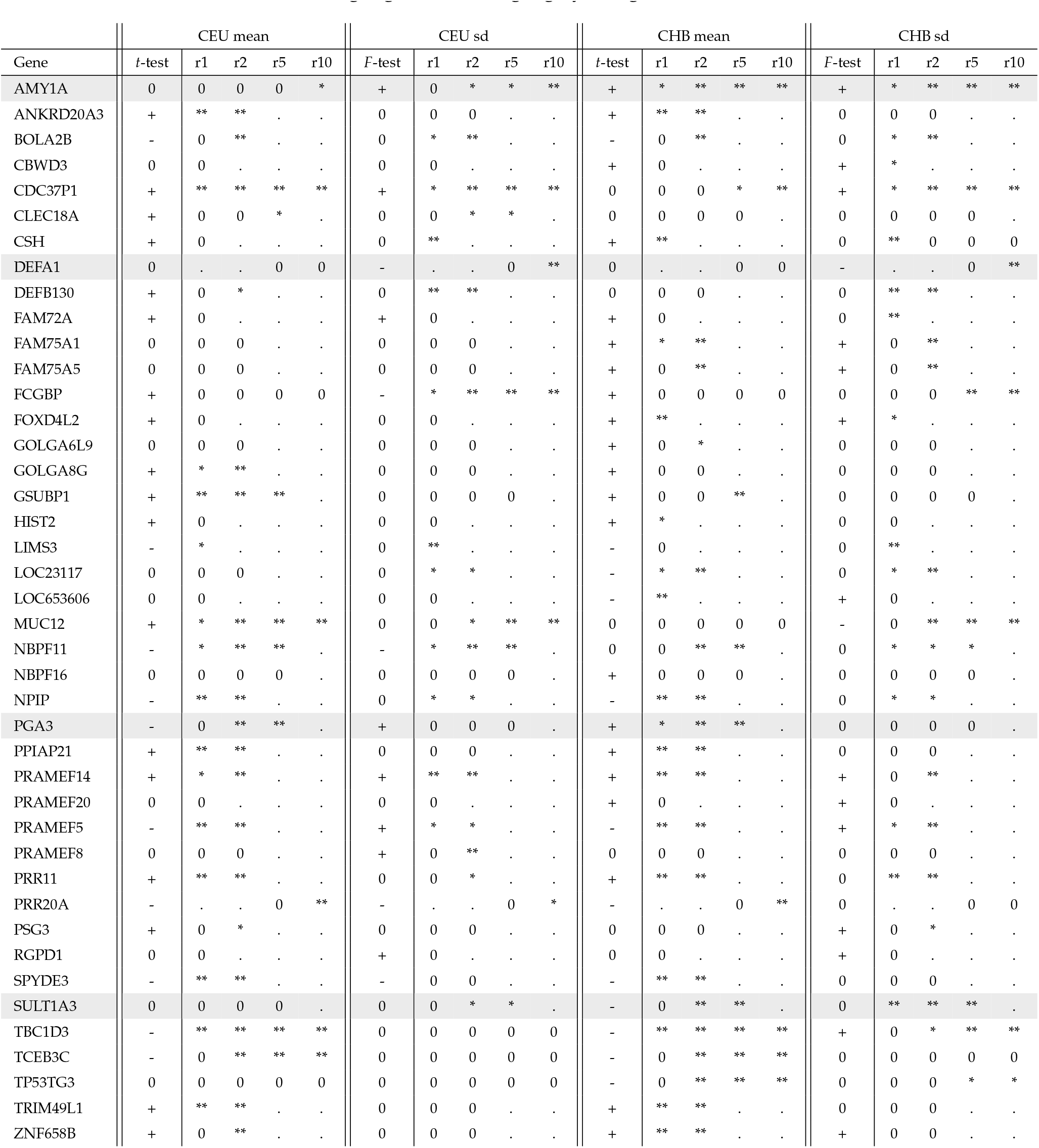
Results of bottleneck simulations. We ran simulations of scenario I (the strongest bottleneck with a reduction to *N* = 100) with parameters (*r, s*_*x*_, *s*_*y*_) estimated from the YRI-data and tested, whether after 1000 generations of recovery the mean and standard deviation *σ* of the CEU and CHB-data can be explained by a bottleneck. Blank space indicates that this parameter combination led to an *s*_*x*_ value out of the range of 0.001 and 0.1, and hence no simulation was run. The columns with 0,+ and – indicate whether there is a significant difference to the empirical data set (see Table 1). In the r1 - r10 columns, a 0 indicates that the data can be explained by a bottleneck. ^*^ and ^**^ show significant differences (5% and 1%) of the simulated and empirical data. The four candidate genes which were used for further simulations are highlighted with a light gray background.

| Gene | CEU mean |  |  |  |  | CEU sd |  |  |  |  | CHB mean |  |  |  |  | CHB sd |  |  |  |  |
| --- | --- | --- | --- | --- | --- | --- | --- | --- | --- | --- | --- | --- | --- | --- | --- | --- | --- | --- | --- | --- |
|  | *t*-test | r1 | r2 | r5 | r10 | *F*-test | r1 | r2 | r5 | r10 | *t*-test | r1 | r2 | r5 | r10 | *F*-test | r1 | r2 | r5 | r10 |
| AMY1A | 0 | 0 | 0 | 0 | * | + | 0 | * | * | ** | + | * | ** | ** | ** | + | * | ** | ** | ** |
| ANKRD20A3 | + | ** | ** | . | . | 0 | 0 | 0 | . | . | + | ** | ** | . | . | 0 | 0 | 0 | . | . |
| BOLA2B | - | 0 | ** | . | . | 0 | * | ** | . | . | - | 0 | ** | . | . | 0 | * | ** | . | . |
| CBWD3 | 0 | 0 | . | . | . | 0 | 0 | . | . | . | + | 0 | . | . | . | + | * | . | . | . |
| CDC37P1 | + | ** | ** | ** | ** | + | * | ** | ** | ** | 0 | 0 | 0 | * | ** | + | * | ** | ** | ** |
| CLEC18A | + | 0 | 0 | * | . | 0 | 0 | * | * | . | 0 | 0 | 0 | 0 | . | 0 | 0 | 0 | 0 | . |
| CSH | + | 0 | . | . | . | 0 | ** | . | . | . | + | ** | . | . | . | 0 | ** | 0 | 0 | 0 |
| DEFA1 | 0 | . | . | 0 | 0 | - | . | . | 0 | ** | 0 | . | . | 0 | 0 | - | . | . | 0 | ** |
| DEFB130 | + | 0 | * | . | . | 0 | ** | ** | . | . | 0 | 0 | 0 | . | . | 0 | ** | ** | . | . |
| FAM72A | + | 0 | . | . | . | + | 0 | . | . | . | + | 0 | . | . | . | 0 | ** | . | . | . |
| FAM75A1 | 0 | 0 | 0 | . | . | 0 | 0 | 0 | . | . | + | * | ** | . | . | + | 0 | ** | . | . |
| FAM75A5 | 0 | 0 | 0 | . | . | 0 | 0 | 0 | . | . | + | 0 | ** | . | . | + | 0 | ** | . | . |
| FCGBP | + | 0 | 0 | 0 | 0 | - | * | ** | ** | ** | + | 0 | 0 | 0 | 0 | 0 | 0 | 0 | ** | ** |
| FOXD4L2 | + | 0 | . | . | . | 0 | 0 | . | . | . | + | ** | . | . | . | + | * | . | . | . |
| GOLGA6L9 | 0 | 0 | 0 | . | . | 0 | 0 | 0 | . | . | + | 0 | * | . | . | 0 | 0 | 0 | . | . |
| GOLGA8G | + | * | ** | . | . | 0 | 0 | 0 | . | . | + | 0 | 0 | . | . | 0 | 0 | 0 | . | . |
| GSUBP1 | + | ** | ** | ** | . | 0 | 0 | 0 | 0 | . | + | 0 | 0 | ** | . | 0 | 0 | 0 | 0 | . |
| HIST2 | + | 0 | . | . | . | 0 | 0 | . | . | . | + | * | . | . | . | 0 | 0 | . | . | . |
| LIMS3 | - | * | . | . | . | 0 | ** | . | . | . | - | 0 | . | . | . | 0 | ** | . | . | . |
| LOC23117 | 0 | 0 | 0 | . | . | 0 | * | * | . | . | - | * | ** | . | . | 0 | * | ** | . | . |
| LOC653606 | 0 | 0 | . | . | . | 0 | 0 | . | . | . | - | ** | . | . | . | + | 0 | . | . | . |
| MUC12 | + | * | ** | ** | ** | 0 | 0 | * | ** | ** | 0 | 0 | 0 | 0 | 0 | - | 0 | ** | ** | ** |
| NBPF11 | - | * | ** | ** | . | - | * | ** | ** | . | 0 | 0 | ** | ** | . | 0 | * | * | * | . |
| NBPF16 | 0 | 0 | 0 | 0 | . | 0 | 0 | 0 | 0 | . | + | 0 | 0 | 0 | . | 0 | 0 | 0 | 0 | . |
| NPIP | - | ** | ** | . | . | 0 | * | * | . | . | - | ** | ** | . | . | 0 | * | * | . | . |
| PGA3 | - | 0 | ** | ** | . | + | 0 | 0 | 0 | . | + | * | ** | ** | . | 0 | 0 | 0 | 0 | . |
| PPIAP21 | + | ** | ** | . | . | 0 | 0 | 0 | . | . | + | ** | ** | . | . | 0 | 0 | 0 | . | . |
| PRAMEF14 | + | * | ** | . | . | + | ** | ** | . | . | + | ** | ** | . | . | + | 0 | ** | . | . |
| PRAMEF20 | 0 | 0 | . | . | . | 0 | 0 | . | . | . | + | 0 | . | . | . | + | 0 | . | . | . |
| PRAMEF5 | - | ** | ** | . | . | + | * | * | . | . | - | ** | ** | . | . | + | * | ** | . | . |
| PRAMEF8 | 0 | 0 | 0 | . | . | + | 0 | ** | . | . | 0 | 0 | 0 | . | . | 0 | 0 | 0 | . | . |
| PRR11 | + | ** | ** | . | . | 0 | 0 | * | . | . | + | ** | ** | . | . | 0 | ** | ** | . | . |
| PRR20A | - | . | . | 0 | ** | - | . | . | 0 | * | - | . | . | 0 | ** | 0 | . | . | 0 | 0 |
| PSG3 | + | 0 | * | . | . | 0 | 0 | 0 | . | . | 0 | 0 | 0 | . | . | + | 0 | * | . | . |
| RGPD1 | 0 | 0 | . | . | . | + | 0 | . | . | . | 0 | 0 | . | . | . | + | 0 | . | . | . |
| SPYDE3 | - | ** | ** | . | . | - | 0 | 0 | . | . | - | ** | ** | . | . | 0 | 0 | 0 | . | . |
| SULT1A3 | 0 | 0 | 0 | 0 | . | 0 | 0 | * | * | . | - | 0 | ** | ** | . | 0 | ** | ** | ** | . |
| TBC1D3 | - | ** | ** | ** | ** | 0 | 0 | 0 | 0 | 0 | - | ** | ** | ** | ** | + | 0 | * | ** | ** |
| TCEB3C | - | 0 | ** | ** | ** | 0 | 0 | 0 | 0 | 0 | - | 0 | ** | ** | ** | 0 | 0 | 0 | 0 | 0 |
| TP53TG3 | 0 | 0 | 0 | 0 | 0 | 0 | 0 | 0 | 0 | 0 | - | 0 | ** | ** | ** | 0 | 0 | 0 | * | * |
| TRIM49L1 | + | ** | ** | . | . | 0 | 0 | 0 | . | . | + | ** | ** | . | . | 0 | 0 | 0 | . | . |
| ZNF658B | + | 0 | ** | . | . | 0 | 0 | 0 | . | . | + | ** | ** | . | . | + | 0 | 0 | . | . |

If we consider a significant difference in the mean of CEU compared to YRI (28 genes; first column in Table 4 to be non-zero), we find that only 17 out of 65 simulated parameter combinations in scenario I can explain these differences. For significant mean changes in CHB (33 genes; Table 4), 22 out of 72 parameter combinations are compatible with the observation, while the remaining 50 can not explain the significant difference. For other examples, `AMY1A` and `PGA3` both showed an increased mean value in CHB. In neither case, and for neither parameter combination, scenario I is sufficient to explain the observation.

From the candidates with a significant difference in mean or variance we selected the three genes coding for digestive enzymes, `AMY1A, SULT1A3, PGA3`, and the defense gene `DEFA1` for a more detailed analysis and tested the GADMA model (Noskova et al. 2020) without and with selection change according to the estimates from regression (scenario IV-VI).

Fig 5 shows mean copy number and coefficient of variation 𝒞_*V*_ at present, simulated according to scenarios IV and VI for 10,000 replicates each. As in scenarios I-III, the terminal values generated in scenario IV are close to the initial ones of the ancestral YRI data set (black line). Therefore, even the more realistic GADMA migration model often fails to explain the data found in CEU and CHB when considering constant selection parameters derived from the ancestral YRI population.

**Figure 5.**
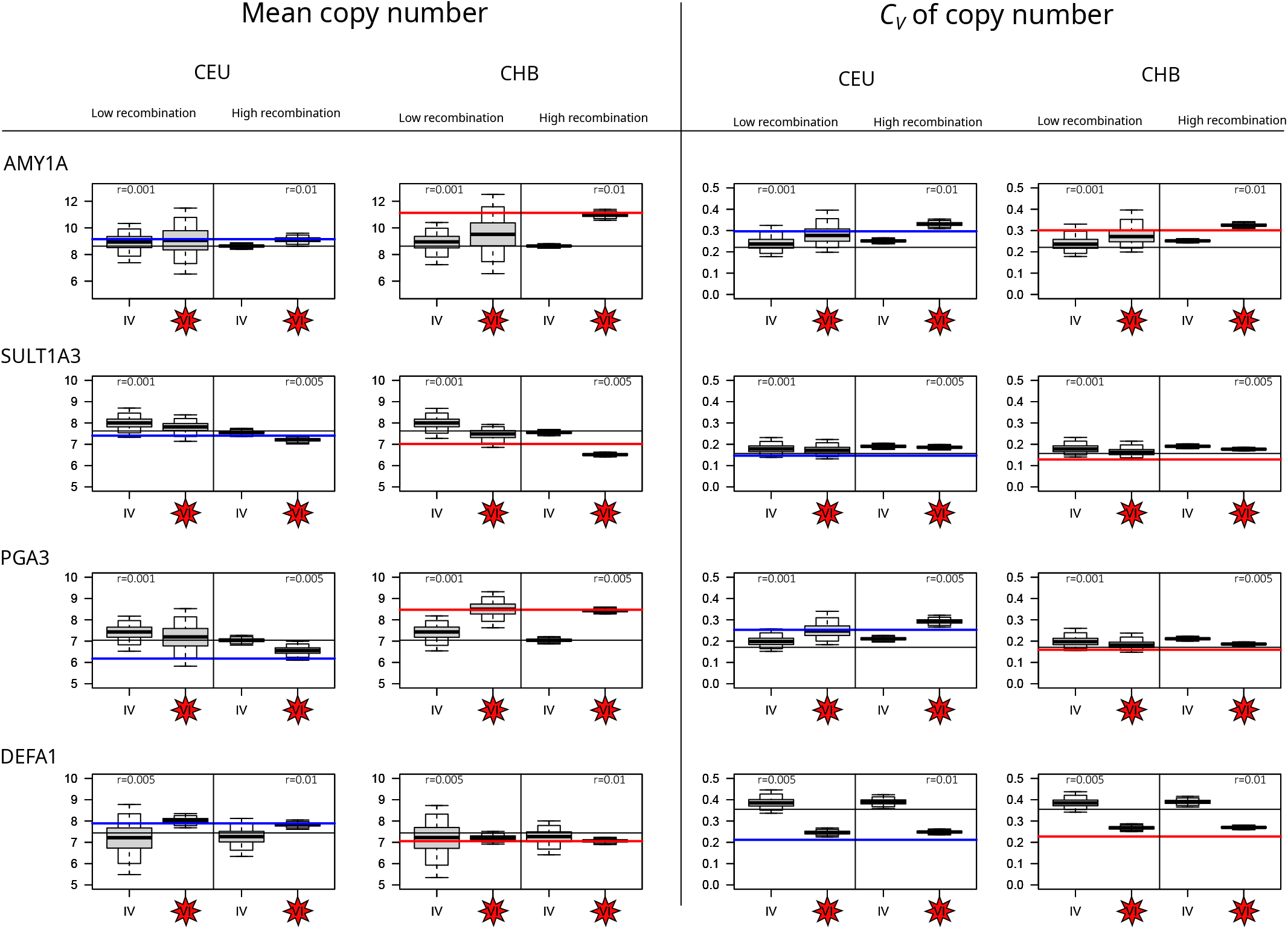
Mean copy number 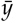, and coefficient of variation 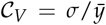 for four candidate genes (`AMY1A, SULT1A3, PGA3, DEFA1`). Box-plots: simulation results for the GADMA demographic model without (scenario IV) and with (scenario VI) change of selection and for two settings of recombination rate (low and high, see Table 3). Horizontal lines: mean and 𝒞_*V*_ of the experimental data in YRI (black), CEU (blue) and CHB (red).

However, when selection strength is allowed to change, as in scenario VI, a different picture emerges: Consider a change of *s*_*x*_ and *s*_*y*_ at 500 generations before present with respect to the values estimated from equation (3), given in Table 3. Then, the simulations return mean and 𝒞_*V*_ which are closer to the values found in CEU and CHB data. Indeed, the empirical data often lie within the 95%- or 99%-quantiles of the simulated data distributions. We observe no strong difference between the results of scenarios V and VI, suggesting that even 500 generations represent a sufficiently large time span to reach a new equilibrium.

Hence, one possible explanation in the shift of copy number distribution of the four candidate genes is a change of selection pressure and adaptation.

The `AMY1A` gene, which encodes amylase, an enzyme that breaks down starch, has strongly increased mean and *σ* in the Asian population, likely linked to adaptations to high grain intake. In the European population, while the variation is increased, the change in mean copy number is small. These findings are in agreement with results of several studies that indicate that individuals from populations with high-starch diets have, on average, more gene copies than those with traditionally low-starch diets (Perry *et al*. 2007; Pajic et al. 2019; Atkinson et al. 2018). Under our model selection strength is relaxed in CEU and CHB with a factor of 4, such that a higher copy number is not selected against and a more widespread distribution of CNV can evolve. A recent study (Inchley et al. 2016) has suggested a more complicated model of Amylase evolution, involving two steps: an expansion from one to several copies after the human-Neanderthal split, but before separation of modern human populations, and a subsequent shift of the optimal gene copy number, independently in different populations. This study also suggests that increase of `AMY1` copy number occurred in South America even more dramatically than in East Asia, a hypothesis which should be tested in the framework of our model as soon as suitable data become available.

`SULT1A3` is a gene in the SULT (sulfotransferases) family, which catalyze sulfation of a variety of substrates, especially catecholamines including dopamine and epinephrine (Brix et al. 1999; Dajani et al. 1999). Polymorphisms in `SULT1A3` and `SULT1A4` have been shown to affect metabolism of therapeutic drugs (eg., (Hui and Liu 2015; Bairam et al. 2019)), and these genes have therefore been studied extensively in the framework of medico- and pharmacogenetics (Thomae et al. 2004; Hildebrandt et al. 2004). In the dataset analyzed, it has reduced mean copy number in Asia but not in Europe. The reduced mean (from 7.6 in YRI to 7.0 in CHB) is a significant difference, which can not be explained with a simple bottleneck scenario with recombination rate higher than 0.002 (see Table 4). If one considers a change of selection as in scenarios V and VI, we expect a stronger selective pressure (rising from *s*_*x*_ = 0.03 to *s*_*x*_ = 0.05 for *r* = 0.002) in CHB. There have been a few studies on copy numbers of SULT1A3/4 genes. Hildebrandt *et al*. (2004) first noted possible duplication of `SULT1A3` and identified a duplicated copy in all four different human populations. More recently, a study of 172 human individuals discovered variable `SULT1A3/4` copy numbers from 1 to 10, and associated its copy number with risk and onset of neurodegenerative disease (Butcher et al. 2017). Note that `SULT1A3` and `SULT1A4` are closely related paralogs that are often difficult to distinguish, and studies on copy number usually put them together.

`PGA3` (Pepsinogen, precursor for pepsin, an enzyme that breaks down protein to smaller peptides) is associated with prostate-specific antigen production. It is the only gene in our list to have opposite changes in two derived populations: its mean copy number increases in Asia and decreases in Europe. As Asian and European humans share most of the same bottleneck period, the diverging copy number distribution is highly unlikely to be a demographic effect, and complex selection patterns are needed to explain the data. Indeed, the bottleneck simulations shown in Table 4 and the simulations V and VI with a change of selection parameters as shown in Fig 5 support this hypothesis. When considering the estimates of Table 3 we observe a small increase of *s*_*x*_ in Asia compared to Africa but a strong decrease of both *s*_*x*_ and *s*_*y*_ in Europe to cope the increased variance in copy number in CEU. The copy number variation on the Pepsinogen (PGA) locus was originally discoverd with electrophoresis and three individual genes (named `PGA 3, 4, 5`) were initially found (Taggart *et al*. 1985). Pepsinogen genes have been shown to duplicate and become lost recurrently in vertebrates (Castro *et al*. 2014). The pepsinogen genes were also shown to have variable expression level in tumor cells, particularly a reduction of PGA expression in esphagael, stomach and thyroid cancers (Shen *et al*. 2020). This could be an additional source of selective pressure besides protein metabolism. While the simplest explanation is that dietary differences between Asian and European populations during the spread of of agriculture (in the last 5000-10000 years) is the driver of `PGA` copy number changes, alternative hypotheses involving tumor suppression or interaction with other enzymes must be considered.

Finally, we analyzed the immune gene Alpha-defensin `DEFA1.` It codes for defensins, proteins that are involved in innate (non-learned) immunity, specifically in antimicrobial defense against a broad spectrum of microorganisms, including bacteria, fungi, and viruses. `DEFA1` shows a decrease in variance in both Asia and Europe, indicating stronger selective pressures. More precisely, when considering the distribution in Fig 1, one observes four individuals in YRI population with high copy number which indicates a relaxed selective pressure in Africa. With equation (3) we find selection coefficients 10-fold smaller in Africa than in Europe and Asia (see Table 3). Alpha-defensins are expressed in neutrophil cells and intestinal epithelial cells, acting as microbiocidal agents (Ganz *et al*. 1985; Ayabe *et al*. 2000; Nassar *et al*. 2007). The genes DEFA1 and `DEFA3` code for some of the Alpha-defensins (HNP1/2/3), and appear to be “interchangeable variant cassettes” within a tandem array of 19kb (Aldred *et al*. 2005). Copy number variation of DEFA1 is present in all apes including gibbon, but the version identified as `DEFA3` is human-specific; the copy number is also demonstrated to affect expression level (Aldred *et al*. 2005). Low copy number of `DEFA1/3` is shown to be associated with hospital-acquired infection (Zhao *et al*. 2018) as well as kidney diseases (Ai *et al*. 2016). On the other hand and counterintuitively, a high copy number of `DEFA1/3` may lead to more severe cases of sepsis (Chen *et al*. 2010, 2019) and is associated with Crohn’s Disease (Jespersgaard *et al*. 2011), and thus selected against. The trade-off between infective and autoimmune diseases could lead to selection towards an intermediate copy number of Alpha-defensins. Therefore, our results suggest a possibility that the out-of-Africa expansion is accompanied by such a change in environmental pathogen diversity that a delicately tuned dosage of defensin is required. This can be corroborated by the fact that YRI has a few individuals with very high (outliers) copy numbers of `DEFA1`, which can not be found in CHB or CEU.

In conclusion, while both demographic effects and shifts in selection schemes can result in changes in copy number distributions, in some of our candidate genes the former is not sufficient to explain the observation. Adaptive processes can induce new relationships between copy number and fitness, and impact the resulting copy number distribution. Importantly, changes in the strength, or direction of selection may become manifest not only in mean copy number, but also in the variance or compound statistics, such as the coefficient of variation.

## Data Availability Statement

All data and costum codes used for simulations and analysis can be found at https://github.com/y-zheng/Distinguishing-roles-adaptation-demography-gene-copy-number-changes-human-populations

## Funding

This work has been funded by grants from the German Research Foundation (DFG SFB-1211, subproject B6) to TW.

## Conflict of interest

The authors declare no conflict of interest.

## Notes

### Competing Interest Statement

The authors have declared no competing interest.

### Summary of Updates

Figure 3 and Figure 5 together with their legends have been updated.

## Literature cited

Ai Z, Li M, Liu W, Foo JN, Mansouri O, Yin P, Zhou Q, Tang X, Dong X, Feng S et al. 2016. Low α-defensin gene copy number increases the risk for IgA nephropathy and renal dysfunction. Science Translational Medicine. 8.

Aldred PM, Hollox EJ, Armour JA. 2005. Copy number polymorphism and expression level variation of the human α-defensin genes DEFA1 and DEFA3. Human Molecular Genetics. 14:2045–2052.

Atkinson FS, Hancock D, Petocz P, Brand-Miller JC. 2018. The physiologic and phenotypic significance of variation in human amylase gene copy number. The American Journal of Clinical Nutrition. 108:737–748.

Ayabe T, Satchell DP, Wilson CL, Parks WC, Selsted ME, Ouellette AJ. 2000. Secretion of microbicidal α-defensins by intestinal paneth cells in response to bacteria. Nature Immunology. 1:113–118.

Bairam AF, Rasool MI, Alherz FA, Abunnaja MS, Daibani AAE, Gohal SA, Alatwi ES, Kurogi K, Liu MC. 2019. Impact of SULT1a3/SULT1a4 genetic polymorphisms on the sulfation of phenylephrine and salbutamol by human SULT1a3 allozymes. Pharmacogenetics and Genomics. 29:99–105.

Brahmachary M, Guilmatre A, Quilez J, Hasson D, Borel C, Warburton P, Sharp AJ. 2014. Digital genotyping of macrosatellites and multicopy genes reveals novel biological functions associated with copy number variation of large tandem repeats. PLoS Genetics. 10:e1004418.

Brix LA, Barnett AC, Duggleby RG, Leggett B, McManus ME. 1999. Analysis of the substrate specificity of human sulfotransferases SULT1a1 and SULT1a3: site-directed mutagenesis and kinetic studies. Biochemistry. 38:10474–10479.

Butcher NJ,, Horne MK, Mellick GD, Fowler CJ, Masters CL, Minchin RF. 2017. Sulfotransferase 1a3/4 copy number variation is associated with neurodegenerative disease. The Pharmacogenomics Journal. 18:209–214.

Carpenter D, Dhar S, Mitchell LM, Fu B, Tyson J, Shwan NAA, Yang F, Thomas MG, Armour JAL. 2015. Obesity, starch digestion and amylase: association between copy number variants at human salivary (AMY1) and pancreatic (AMY2) amylase genes. Human Molecular Genetics. 24:3472–3480.

Carvalho CMB, Lupski JR. 2016. Mechanisms underlying structural variant formation in genomic disorders. Nature Reviews Genetics. 17:224–238.

Castro LFC, Goncalves O, Mazan S, Tay BH, Venkatesh B, Wilson JM. 2014. Recurrent gene loss correlates with the evolution of stomach phenotypes in gnathostome history. Proceedings of the Royal Society B: Biological Sciences. 281:20132669.

Chen Q, Hakimi M, Wu S, Jin Y, Cheng B, Wang H, Xie G, Ganz T, Linzmeier RM, Fang X. 2010. Increased genomic copy number of DEFA1/DEFA3 is associated with susceptibility to severe sepsis in chinese han population. Anesthesiology. 112:1428–1434.

Chen Q, Yang Y, Hou J, Shu Q, Yin Y, Fu W, Han F, Hou T, Zeng C, Nemeth E et al. 2019. Increased gene copy number of DEFA1/DEFA3 worsens sepsis by inducing endothelial pyroptosis. Proceedings of the National Academy of Sciences. 116:3161–3170.

Dajani R, Sharp S, Graham S, Bethell SS, Cooke RM, Jamieson DJ, Coughtrie MW. 1999. Kinetic properties of human dopamine sulfotransferase (SULT1a3) expressed in prokaryotic and eukaryotic systems: Comparison with the recombinant enzyme purified fromEscherichia coli. Protein Expression and Purification. 16:11–18.

Falchi M, Moustafa JSES, Takousis P, Pesce F, Bonnefond A, Andersson-Assarsson JC, Sudmant PH, Dorajoo R, Al-Shafai MN, Bottolo L et al. 2014. Low copy number of the salivary amylase gene predisposes to obesity. Nature Genetics. 46:492–497.

Fu YX. 1997. Statistical tests of neutrality of mutations against population growth, hitchhiking and background selection. Genetics. 147:915–925.

Ganz T, Selsted ME, Szklarek D, Harwig SS, Daher K, Bainton DF, Lehrer RI. 1985. Defensins. natural peptide antibiotics of human neutrophils. Journal of Clinical Investigation. 76:1427–1435.

Hildebrandt MA, Salavaggione OE, Martin YN, Flynn HC, Jalal S, Wieben ED, Weinshilboum RM. 2004. Human SULT1a3 pharmacogenetics: gene duplication and functional genomic studies. Biochemical and Biophysical Research Communications. 321:870–878.

Hui Y, Liu MC. 2015. Sulfation of ritodrine by the human cytosolic sulfotransferases (SULTs): Effects of SULT1a3 genetic polymorphism. European Journal of Pharmacology. 761:125–129.

Inchley CE, Larbey CDA, Shwan NAA, Pagani L, Saag L, Antao T, Jacobs G, Hudjashov G, Metspalu E, Mitt M et al. 2016. Selective sweep on human amylase genes postdates the split with neanderthals. Scientific Reports. 6.

Iskow RC, Gokcumen O, Lee C. 2012. Exploring the role of copy number variants in human adaptation. Trends in Genetics. 28:245–257.

Jespersgaard C, Fode P, Dybdahl M, Vind I, Nielsen OH, Csillag C, Munkholm P, Vainer B, Riis L, Elkjaer M et al. 2011. Alpha-defensin DEFA1a3 gene copy number elevation in danish crohn’s disease patients. Digestive Diseases and Sciences. 56:3517–3524.

Liao WW, Asri M, Ebler J, Doerr D, Haukness M, Hickey G, Lu S, Lucas JK, Monlong J, Abel HJ et al. 2023. A draft human pangenome reference. Nature. 617:312–324.

Lohmueller KE. 2014. The impact of population demography and selection on the genetic architecture of complex traits. PLoS Genetics. 10:e1004379.

Nassar H, Lavi E, Akkawi S, Bdeir K, Heyman SN, Raghunath P, Tomaszewski J, Higazi AAR. 2007. α-defensin: Link between inflammation and atherosclerosis. Atherosclerosis. 194:452–457.

Noskova E, Ulyantsev V, Koepfli KP, O’Brien SJ, Dobrynin P. 2020. GADMA: Genetic algorithm for inferring demographic history of multiple populations from allele frequency spectrum data. GigaScience. 9.

Otto M, Zheng Y, Wiehe T. 2022. Recombination, selection, and the evolution of tandem gene arrays. Genetics. 221.

Pajic P, Pavlidis P, Dean K, Neznanova L, Romano RA, Garneau D, Daugherity E, Globig A, Ruhl S, Gokcumen O. 2019. Independent amylase gene copy number bursts correlate with dietary preferences in mammals. eLife. 8.

Perry GH, Dominy NJ, Claw KG, Lee AS, Fiegler H, Redon R, Werner J, Villanea FA, Mountain JL, Misra R et al. 2007. Diet and the evolution of human amylase gene copy number variation. Nature Genetics. 39:1256–1260.

Sebat J, Lakshmi B, Troge J, Alexander J, Young J, Lundin P, anér SM, Massa H, Walker M, Chi M et al. 2004. Large-scale copy number polymorphism in the human genome. Science. 305:525–528.

Shen S, Li H, Liu J, Sun L, Yuan Y. 2020. The panoramic picture of pepsinogen gene family with pan-cancer. Cancer Medicine. 9:9064–9080.

Stajich JE. 2004. Disentangling the effects of demography and selection in human history. Molecular Biology and Evolution. 22:63–73.

Sudmant PH, Kitzman JO, Antonacci F, Alkan C, Malig M, Tsalenko A, Sampas N, Bruhn L, Shendure J, and EEE. 2010. Diversity of human copy number variation and multicopy genes. Science. 330:641–646.

Sudmant PH, Mallick S, Nelson BJ, Hormozdiari F, Krumm N, Huddleston J, Coe BP, Baker C, Nordenfelt S, Bamshad M et al. 2015. Global diversity, population stratification, and selection of human copy-number variation. Science. 349.

Taggart RT, Mohandas TK, Shows TB, Bell GI. 1985. Variable numbers of pepsinogen genes are located in the centromeric region of human chromosome 11 and determine the high-frequency electrophoretic polymorphism. Proceedings of the National Academy of Sciences. 82:6240–6244.

Tajima F. 1989. Statistical method for testing the neutral mutation hypothesis by DNA polymorphism. Genetics. 123:585–595.

Thomae BA, Rifki OF, Theobald MA, Eckloff BW, Wieben ED, Weinshilboum RM. 2004. Human catecholamine sulfotransferase (SULT1a3) pharmacogenetics: functional genetic polymorphism. Journal of Neurochemistry. 87:809–819.

Usher CL, Handsaker RE, Esko T, Tuke MA, Weedon MN, Hastie AR, Cao H, Moon JE, Kashin S, Fuchsberger C et al. 2015. Structural forms of the human amylase locus and their relationships to SNPs, haplotypes and obesity. Nature Genetics. 47:921–925.

Zhao J, Gu Q, Wang L, Xu W, Chu L, Wang Y, Li Z, Wu S, Xu J, Hu Z et al. 2018. Low-copy number polymorphism in DEFA1/DEFA3 is associated with susceptibility to hospital-acquired infections in critically ill patients. Mediators of Inflammation. 2018:1–8.

