## Supplementary figures and images for "Detecting adaptive changes in gene copy number distribution accompanying the human out-of-Africa expansion"

### Fig_S1-Regression_1.png

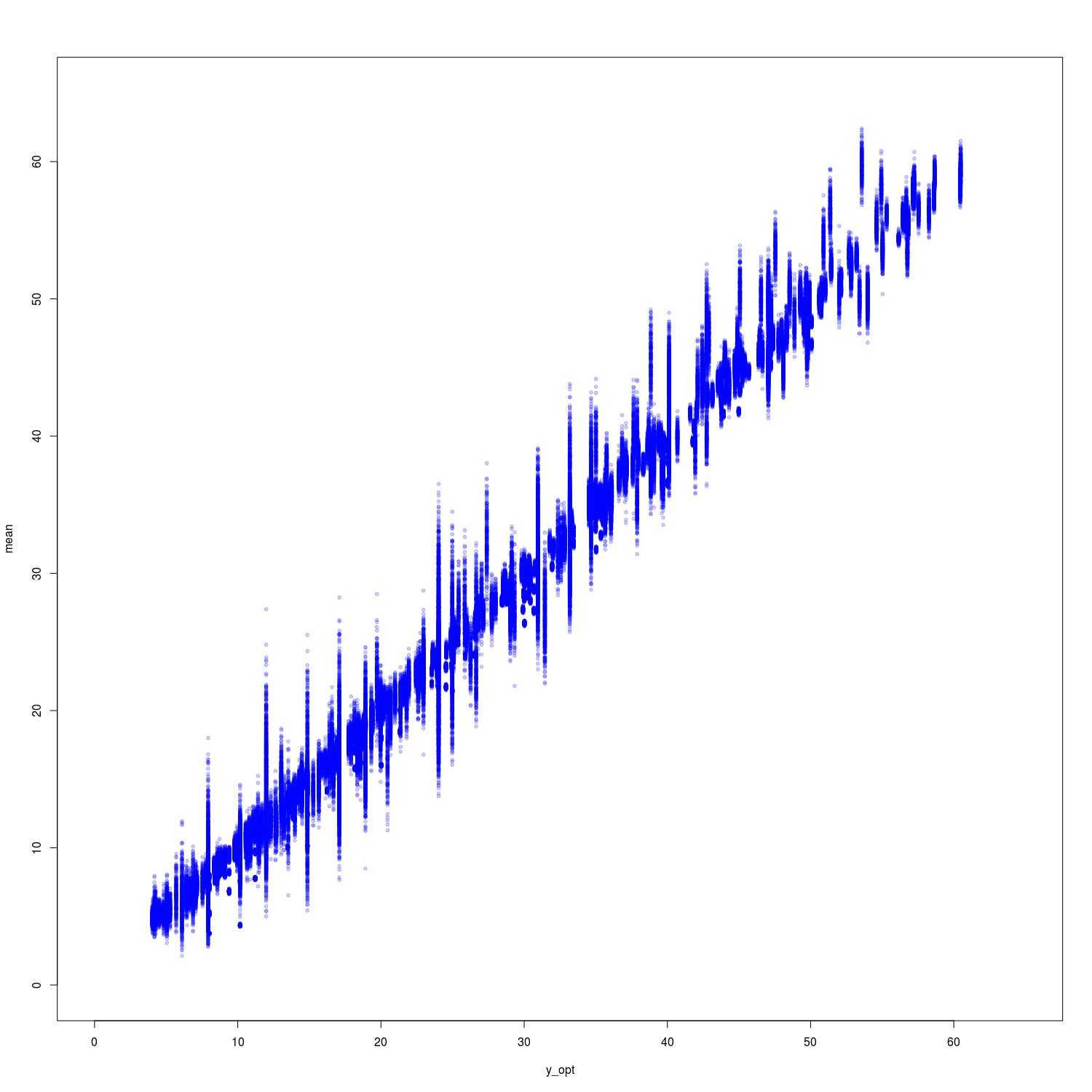

### Fig_S1-Regression_2.png

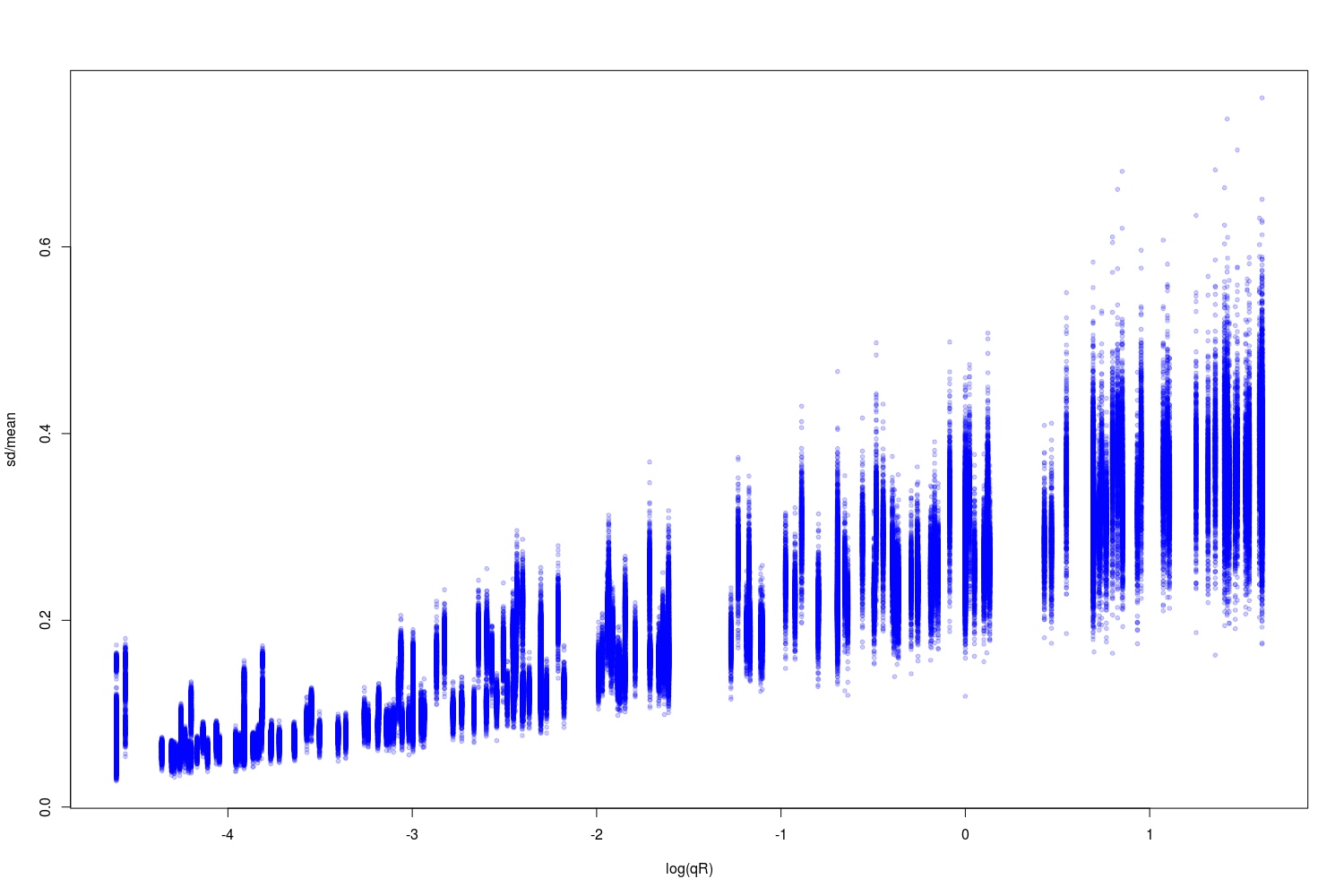
